# Engineering multiple levels of specificity in an RNA viral vector

**DOI:** 10.1101/2020.05.27.119909

**Authors:** Xiaojing J. Gao, Lucy S. Chong, Michaela H. Ince, Matthew S. Kim, Michael B. Elowitz

**Author notes:** equal contribution (Xiaojing J. Gao, Lucy S. Chong).

## Abstract

Synthetic molecular circuits could provide powerful therapeutic capabilities, but delivering them to specific cell types and controlling them remains challenging. An ideal “smart” viral delivery system would enable controlled release of viral vectors from “sender” cells, conditional entry into target cells based on cell-surface proteins, conditional replication specifically in target cells based on their intracellular protein content, and an evolutionarily robust system that allows viral elimination with drugs. Here, combining diverse technologies and components, including pseudotyping, engineered bridge proteins, degrons, and proteases, we demonstrate each of these control modes in a model system based on the rabies virus. This work shows how viral and protein engineering can enable delivery systems with multiple levels of control to maximize therapeutic specificity.

## Introduction

The ability to deliver a designed nucleic acid to target cell types would open up powerful possibilities for basic research and therapeutic applications. Improved delivery capabilities could enable new generations of gene therapy that take advantage of advances in synthetic biology to provide increased specificity and control. In fact, synthetic biologists have now developed a broad range of biological circuit designs that sense and respond to endogenous cellular states^1–3^. These include systems based on protein-DNA interaction^4^, regulatory RNAs^5^, and protein-level circuits^6–8^, as well as combinations of these modalities^9,10^. By operating intracellularly, they would not be limited to sensing surface proteins, but could instead directly interrogate the core pathways that drive cellular behaviors, allowing conditional responses to specific cellular states. While circuit engineering has progressed enormously, our limited ability to deliver circuits to cells in vivo has prevented their therapeutic use. A specific, effective, and controllable method to transfer circuits into target cell types could provide a foundation for the future development of circuits as therapeutics.

Viruses possess many powerful capabilities as delivery systems. Viruses can preferentially infect specific cell types, and then replicate intracellularly to high levels, enabling strong expression of virally encoded proteins, potentially including engineered “cargo” genes. Researchers have therefore engineered diverse classes of DNA and RNA viruses for gene therapy ^11,12^ and cancer therapeutic ^13^ applications. Among these, RNA riboviruses ^11,12,14^ (RNA viruses, excluding retroviruses) offer unique advantages, since they remain at the RNA level, thereby avoiding integration into the host genome and potential mutagenesis of the host. More specifically, viruses in the order *Mononegavirales* have compact, well-studied genomes that offer multiple avenues for engineering control, can support high level expression of protein components, and exhibit relatively high genomic stability ^15^. Notable examples of engineered viruses from this order include vesicular stomatitis virus (VSV) for oncolytic viral therapies ^16^, Sendai virus for stem cell reprogramming ^17^, and rabies virus for synaptic tracing ^18^.

Rabies virus could provide an ideal platform to demonstrate multiple levels of external and internal control with minimal risk of host genome integration. Rabies virus has been well-characterized and extensively used in neurobiology contexts. Further, its ability to spread can be limited and controlled by deleting the glycoprotein, G, from the viral genome and supplying it in the host host cell, in *trans* ^*19*^. Such G-deleted rabies viruses can also be pseudotyped to target specific cell surface proteins ^20^. Most recently, researchers have explored the possibility of making rabies infection transient through modification of an essential viral protein ^21^.

Nevertheless, transforming rabies virus into a practical circuit delivery system requires integrating existing control mechanisms into a single platform, and developing additional capabilities that together allow independent control of multiple stages of the viral life cycle. To limit viral production in space and time, this system should enable control of viral exit from virus-producing cells based on inducers or environmental signals. To restrict the virus to specific targeted cell types, it should also limit viral entry to cells expressing selected cell-surface antigens and make viral replication conditional on the expression of intracellular proteins. For external control, the overall system should be controllable with a well characterized and safe small-molecule drug. To maximize safety and minimize unintended toxicity from the virus itself, viral infection should be reversible through a control system that is robust to mutations.

Here, we set out to develop an integrated system that enables control of viral release, entry, and replication by combining existing rabies virus control mechanisms with new mechanisms for control of viral replication (Fig. 1). These results demonstrate and integrate multiple levels of control, and provide a framework for engineering a broadly useful RNA viral delivery platform for emerging applications in therapeutic synthetic biology and conventional gene therapy.

**Figure 1.**
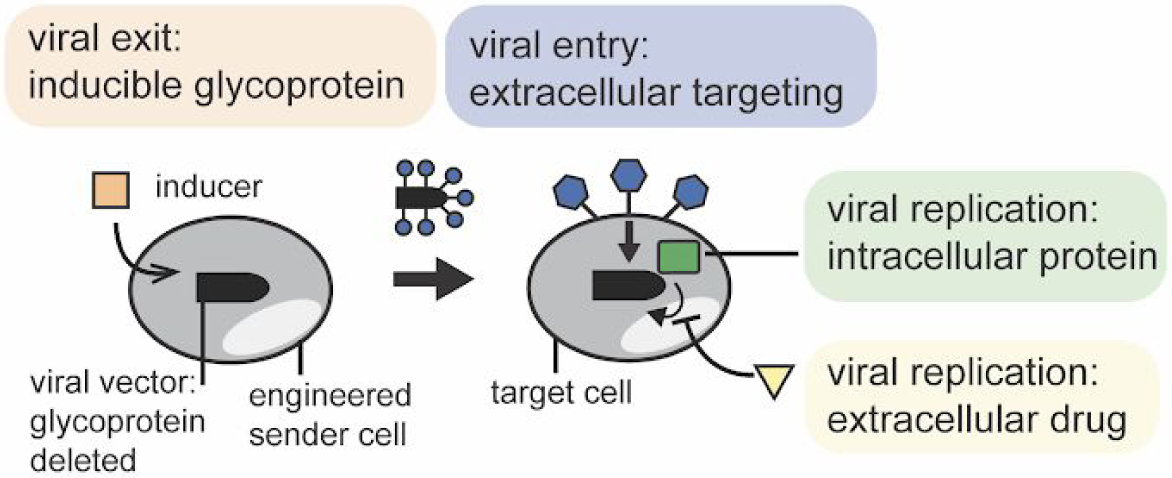
An ideal viral circuit delivery system with four distinct levels of control. Sender cells (left) would be engineered to secrete the virus under the control of an inducer (orange square). Produced viruses would be packaged using a heterologous glycoprotein (“pseudotyped”, blue circles) to selectively infect target cells (right) expressing a desired surface antigen (blue hexagons). Once inside the target cell, viral replication would be conditional on the presence of an intracellular protein (green square). For safety, it would also be possible to suppress viral replication and eliminate the virus using a drug (yellow triangle).

## Results

### Doxycycline-inducible expression of glycoprotein controls viral exit from sender cells

We first engineered a controllable ‘sender’ cell line that releases viral particles in response to external induction. The system takes advantage of the well-characterized paradigm of glycoprotein (G) trans-complementation, in which the G gene can be removed from the viral genome and supplied instead in the host cell line, in order to permit single step infection ^18^ (Fig. 2a). We deleted the native viral G gene, replacing it with mCherry for visualization (RVdG). We then incorporated a single copy of the G transgene in the host genome using the Flp-In system in HEK293 cells (Fig. 2a-b, Methods). The ectopic G gene was controlled by a CMV-TO promoter that was normally repressed by constitutively expressed TetR, but could be readily induced by addition of doxycycline (dox) (Fig. 2a-b, Methods). We also stably incorporated a nuclear-localized Citrine (H2B-Citrine) as a marker. This design can be adapted to allow conditional regulation of G protein expression by other transcriptional inputs, including natural or synthetic signaling pathways ^22–24^.

**Figure 2.**
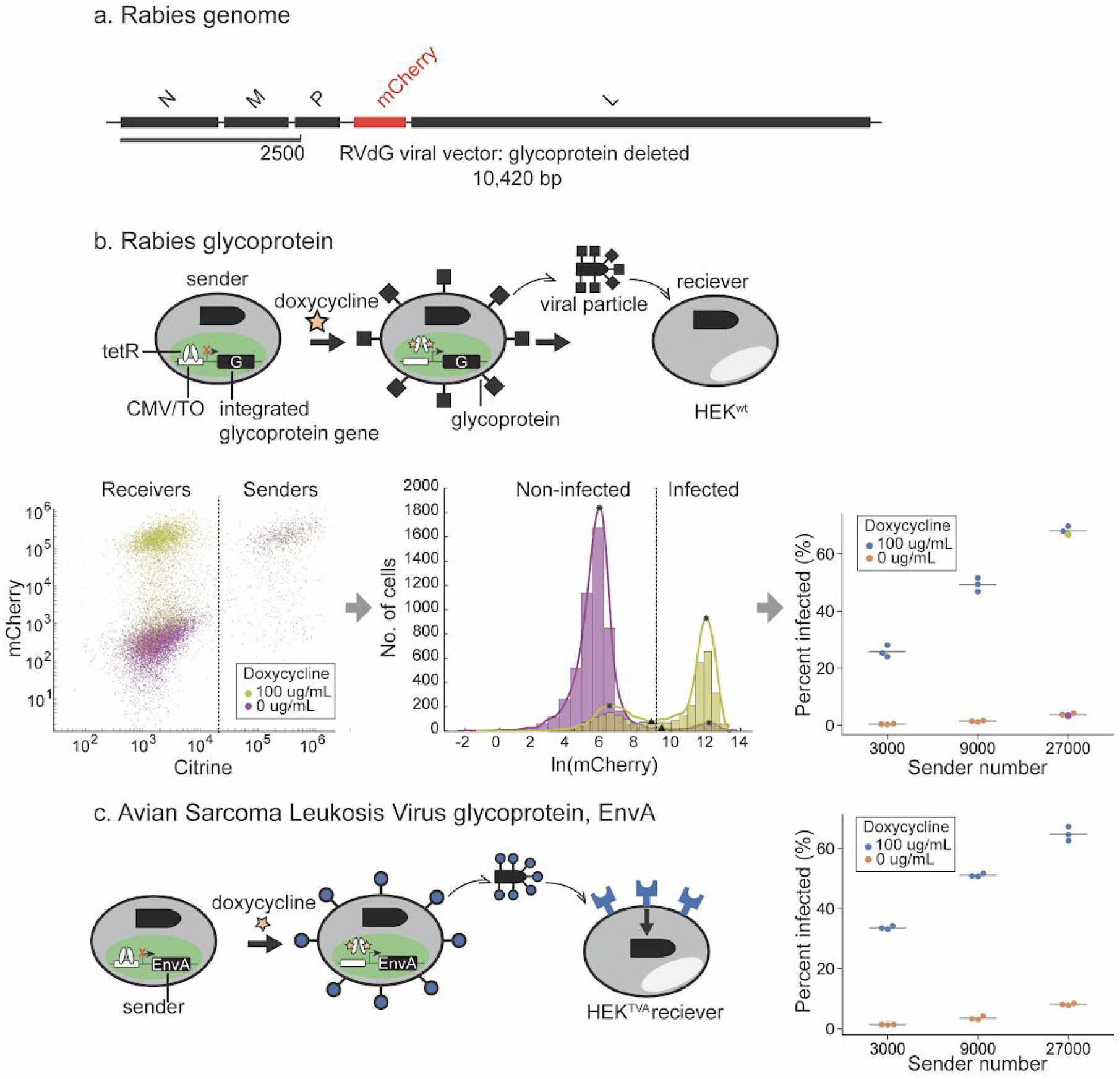
Doxycycline-inducible expression of glycoprotein controls viral exit from sender cells. **(a)** Of the five proteins encoded by the rabies viral genome, the glycoprotein (G) gene is replaced with mCherry (RVdG). The RVdG virus is reconstituted and propagated in producer cells that provide G in trans from the cellular genome. **(b)** Top, sender cell lines with a single copy of genomically integrated, doxycycline-inducible G for the controlled secretion of RVdG. The senders are also labeled with constitutively expressed Citrine. Wildtype HEK293 (HEK^wt^) target cells are co-cultured with the senders to quantify the release of infection competent RVdG. Left, flow cytometry of sender and receiver co-culture in the presence or absence of doxycycline. Vertical line separates receiver and sender subpopulations. Middle, distribution of mCherry signal in HEK^wt^ in the presence or absence of doxycycline (after gating out Citrine positive sender cells) measured with flow cytometry. Vertical line indicates mean local minima used to threshold between infected and weakly infected/non-infected cells. The “percent infected” metric reflects the fraction of cells with signal above the threshold (see Methods). The data points in the swarmplot corresponding to the examples are indicated with stars. Similar swarmplots are used in subsequent panels. Right, percent of target cell infected in the presence or absence of doxycycline under varying numbers of sender cells. **(c)** Replacing the G protein in **b** with a sarcoma leukosis virus glycoprotein gene (EnvA) under a doxycycline-inducible promoter. Right, percent of TVA-displaying target cell (HEK^TVA^) infected in the presence or absence of doxycycline under varying numbers of sender cells.

To validate the inducibility of viral release, we co-cultured a minority of RVdG-infected sender cells with a majority of HEK293 target cells (HEK^wt^), in the presence or absence of doxycycline. After 3 days, we measured the level of mCherry signal in target cells using flow cytometry. Target cell mCherry intensities exhibited a bimodal distribution, consistent with the presence of uninfected and infected cell populations (Fig. 2b). Further, the infected (high mCherry intensity) sub-population showed relatively uniform (coefficient of variation = 0.23 ± 0.01 for triplicates in the “27,000 sender” condition) mCherry expression, suggesting that viral replication drives its concentration to a well-defined concentration within the cell.

Using this system, we next asked how the rate of target cell infection depended on doxycycline induction and sender cell density. The clean separation between the mCherry distributions allowed quantification of infected and uninfected population sizes in different conditions (Fig. 2b, right, Supplementary Methods). As expected, infection rate depended strongly, though not absolutely, on the presence of doxycycline, with low basal levels of infection likely due to leaky expression of G from its CMV-based TetR-repressible promoter (Fig. 2b, right). Infection also exhibited a dose-dependent, but sub-linear, increase with sender cell fraction (Fig. 2b, right).

This approach can be extended to enable controlled secretion of pseudotyped viruses, in which the native viral glycoprotein is replaced with a distinct viral glycoprotein conferring different tropism ^25^. For example, the well-characterized EnvA glycoprotein from the avian sarcoma leukosis virus binds specifically to the avian TVA protein, and abolishes rabies’ original tropism for mammalian cells ^20,26^. As a demonstration, we engineered an EnvA-pseudotyped sender cell line and a cognate target cell line expressing TVA (Fig. 2c, left). As expected, viral infection was strongly dox-dependent and increased in a dose-dependent manner with sender cell number (Fig. 2c, right). Together, these results indicate that inducible glycoprotein expression enables external regulation of viral secretion for viruses both with wildtype G and pseudotyped with EnvA.

### Pseudotyping and bridge proteins control viral entry

Pseudotyping opens up the possibility of targeting engineered rabies viruses to specific mammalian cell types based on their expression of surface proteins. To achieve this, we took inspiration from previous efforts of targeting viral infection to specific cells based on their surface proteins^27–30^. We engineered a bivalent “bridge” protein, consisting of the TVA extracellular domain fused to a nanobody that recognizes GFP ^31^ (Fig. 3a, left). In this modular scheme, viral entry requires both the bridge protein and expression of GFP on the surface of the target cell.

**Figure 3.**
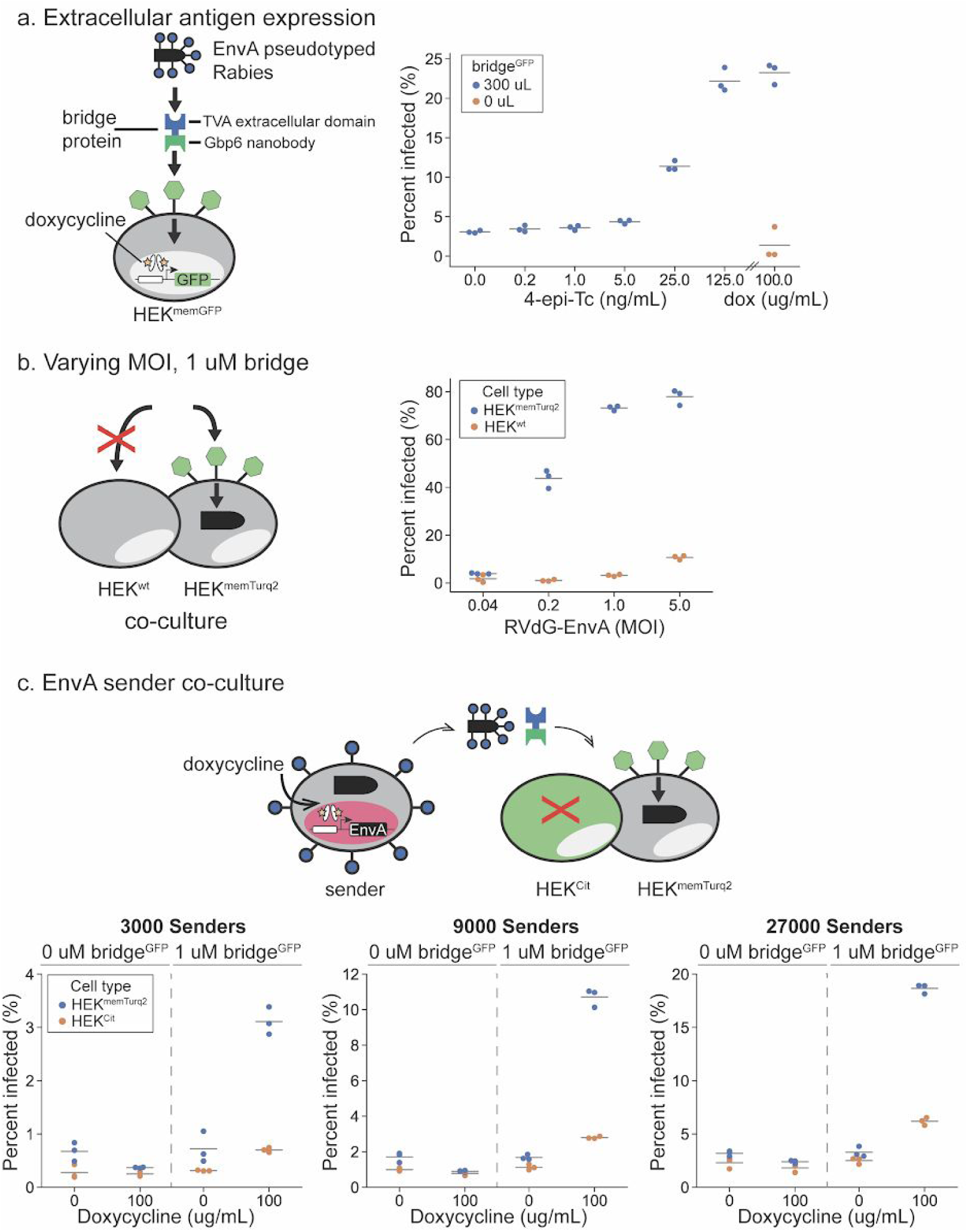
Pseudotyping and bridge proteins control viral entry. **(a)** Left, schematic of the bivalent bridge protein directing EnvA-pseudotyped rabies (RVdG-EnvA) to infect GFP-displaying target cells (HEK^memGFP^). Recombinantly expressed bridge protein contains the TVA receptor extracellular domain and Gbp6 nanobody (bridge^GFP^). Right, percent of HEK^memGFP^ infected under increasing concentrations of 4-epi-Tc using 1 MOI RVdG-EnvA and 300 uL of bridge^GFP^-conditioned media. **(b)** Constant bridge^GFP^ and varying concentrations of RVdG-EnvA administered to a mixed population of HEK^wt^ and HEK^memTurq2^ preferentially infected HEK^memTurq2^ cells. **(c)** Top, co-culture of doxycycline-inducible EnvA sender cells co-expressing IFP with target cells HEK^memGFP^ and non-target cells HEK^Cit^. Bottom, percent of target and non-target cells infected in the presence or absence of doxycycline and bridge protein under varying sender numbers.

We purchased EnvA-pseudotyped rabies virus (RVdG-EnvA), engineered HEK293 derivatives inducibly expressing GFP or Turq2 (which exhibits shifted spectra but is still recognized by the GFP nanobody) on the cell surface as target cells (HEK^memGFP^, HEK^memTurq2^ Fig. 3a, left), and also engineered the bridge protein, termed bridge^GFP^. In order to characterize the specificity of the resulting system, we added RVdG-EnvA to co-cultured parental HEK^wt^ cells and target HEK^memGFP^ cells, either with or without the bridge^GFP^. As expected, infection was strongly enhanced by the combination of surface GFP/Turq2 expression and the addition of bridge^GFP^ (Fig. 3a, right, Supplementary Fig. 1a). Infection of HEK^memGFP^ cells was not detectable without bridge^GFP^ at a multiplicity of infection (MOI) of 1. At much higher MOIs of 5, some non-specific infection of HEK^wt^ and HEK^memGFP^ cells did occur without bridge protein (Fig. 3b). However, the infection rate of HEK^memGFP^ was 8-fold higher with the bridge protein than without it, even at this high MOI value. Furthermore, even at a more modest MOI of ∼1, infection remained dependent on the bridge protein (Supplementary Fig. 1b). The pseudotyping bridge protein strategy thus provided strong infection specificity contingent upon surface proteins. In principle, the same design could be adapted to target natural cell types based on cell surface markers by replacing the GFP-targeting nanobody with a corresponding binding domain.

We next sought to combine inducible secretion with bridge-dependent infection. We co-cultured RVdG infected EnvA sender cells together with both target HEK^memTurq2^ and non-target HEK^Cit^ cells. We then analyzed the relative infection rates of target and non-target cells across different viral secretion rates and numbers of sender cells with or without the bridge protein. As expected, infection strongly depended on the level of induction of the EnvA senders, the presence of bridge^GFP^, and the expression of surface antigen, making infection simultaneously dependent on all three factors (Fig. 3c).

### Viral replication is controlled by an intracellular protein

The ability to control or condition viral replication on intracellular proteins and external inducers would allow more cell type specific control of viral infection. We therefore sought to engineer a replication control system that could be sensitive to the presence of specific intracellular proteins, such as cell type markers or proteins expressed in specific states (e.g. active proliferation), and external small molecule inducers. To connect protein sensing to viral replication in a modular fashion, one needs to design a protein whose activity is required for viral replication but dependent on the presence of an inducer and/or intracellular target protein.

To achieve this, we identified essential viral proteins that can be regulated by the attachment of conditional degradation domains (degrons). The DHFR degradation domain (degron) destabilizes attached proteins, but can be inhibited by the trimethoprim (TMP) ^32^. To identify sensitive sites, we fused DHFR with each essential protein and screened for locations that permitted viral replication in the presence of TMP. The C-terminus of the viral P protein proved to be an ideal site for regulation. It was sensitive to DHFR incorporation, but this effect could be blocked by TMP. We introduced a TEV cleavage site between the P protein and the degron (P-DHFR), so that the virus (RVdG-P-DHFR) can be recovered from a cell line stably expressing the TEV protease without the need for TMP (Supplementary Fig. 3a).

To enable regulation by endogenous proteins, we took advantage of an engineered unstable nanobody that is stabilized by binding to its target antigen^33^. We replaced the DHFR in RVdG-P-DHFR with the destabilized GFP nanobody, denoted GBP (RVdG-P-GBP, Fig. 4a, 4b top). To test whether viral replication was indeed conditional on expression of intracellular GFP, or more precisely its yellow fluorescent variant, Citrine, we used RVdG-P-GBP to infect a co-culture of Citrine-expressing HEK^Cit^ cells and non-expressing parental HEK^wt^ cells (Fig 4b, bottom left). As measured by mCherry expression, the citrine-positive cells were infected at high rates (Fig. 4a, bottom right), while infection levels in citrine-negative cells were indistinguishable from background (Fig. 4a, bottom right). This result indicates that rabies virus replication can be made dependent on the presence of an unrelated intracellular protein (GFP).

**Figure 4.**
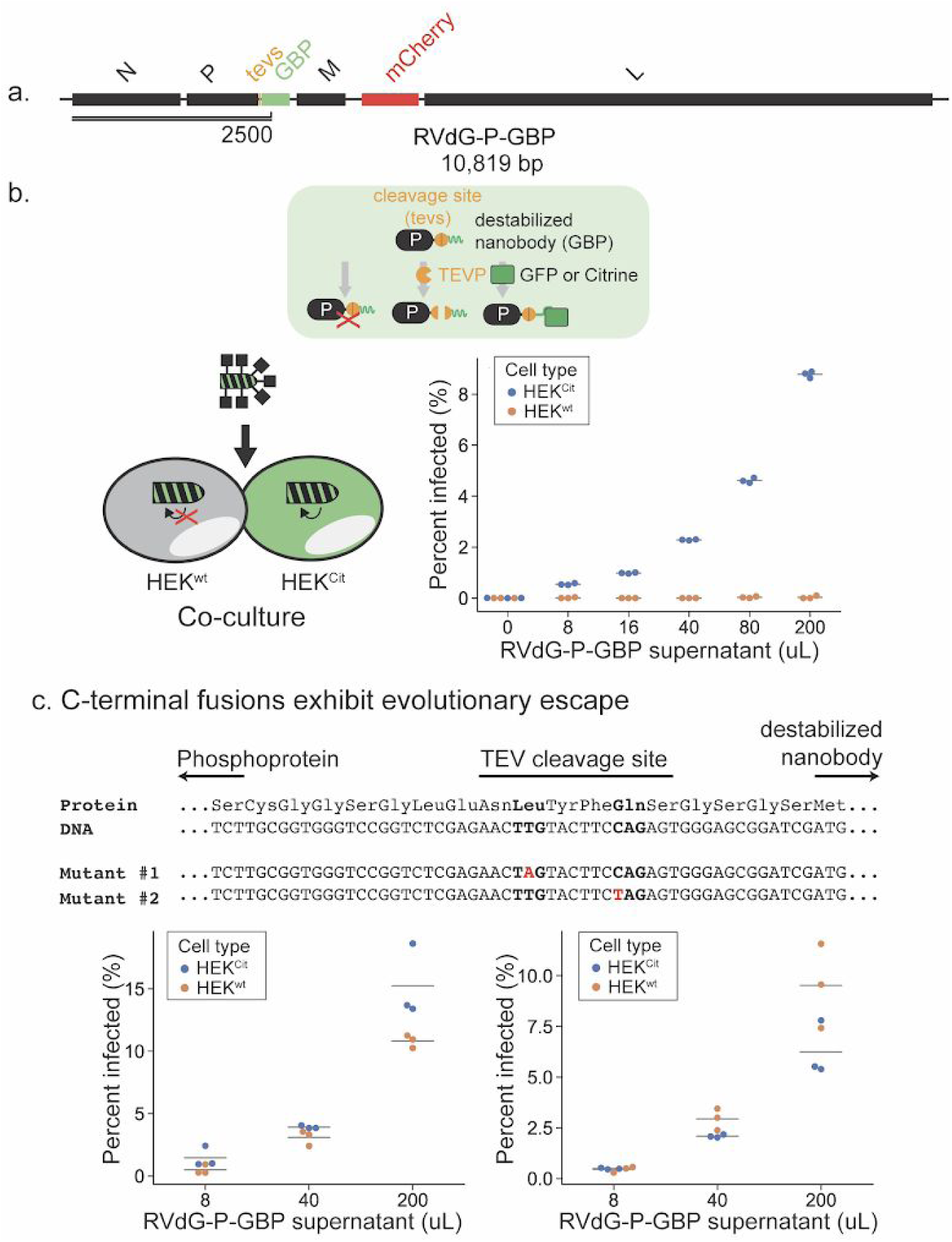
Viral replication is controlled by an intracellular protein. **(a)** Recombinant rabies genome, RVdG-P-GBP. **(b)** Tagging the phosphoprotein (P) with a degron controls viral replication. Top, Schematic of a P protein (black rectangle) tagged with a destabilized nanobody, GBP (green wavy line), and an intervening TEV protease cleavage site (orange circle). P is stabilized when GFP is removed by TEV protease or stabilized when bound to GFP or Citrine, thus permitting viral replication. Bottom left, in a mixed population of HEK^wt^ and HEK293 with cytoplasmic expression of Citrine (HEK^Cit^, see Methods), RVdG-P-GBP will preferentially replicate in HEK^Cit^. Right, increasing concentration of RVdG-P-GBP infects HEK^Cit^, but not HEK^wt^. **(c)** Top, the P-GBP design exhibits evolutionary escape. Sanger sequencing of the junction region between P and GBP of two escape mutants identified two distinct nonsense mutations. Red indicates single nucleotide mutations and bold indicates mutated codons. **Bottom**, infection of a mixed population of HEK^wt^ and HEK^Cit^ using mutant RVdG-PGBP showed diminished discrimination.

Despite the success of the P-degron strategy, conditionality was diminished after ∼1 month of continuous viral passaging in the producer cells (Fig. 4c, bottom, Supplementary Table 2, see Methods). Given the error rate of rabies RNA-dependent RNA polymerase^34^, we reasoned that this loss of function could arise from mutations that rescue P function in the absence of the target protein. Indeed, when we sequenced the P-degron junction in the RVdG-P-GBP viral genome, we consistently found nonsense mutations that truncate the protein before the degron is translated (Fig. 4c, top, Supplementary Fig. 3c), bypassing regulation. Similar “escape mutants” have been independently reported in other work ^34^, indicating that engineering evolutionary robustness is essential.

### An external drug inhibits viral replication and permits viral removal

To reduce selection pressure for escape mutants and improve evolutionary robustness in our intended use case (drug-controlled reversal of established infection) we inserted an HCV protease (HCVP) flanked by its cognate cleavage sites ^35–38^ between P and L (RVdG-P-HCVP-L, Fig. 5a). In this design, HCVP normally acts to cleave both sites, separating the P and L proteins and allowing them to function (Fig. 5b, left). Critically, however, the drug asunaprevir (ASV) blocks HCV protease activity ^39^, leaving P and L tethered unproductively together, disrupting viral replication (Fig. 5b, left). This strategy disfavors selection for escaper mutants in two ways. First, contrary to the previous strategy, here ASV is absent when the virus is being passaged or infection is being established (see the “curing” experiment below), relieving the selection pressure for HCVP to lose sensitivity to ASV inhibition. Second, point mutations or deletions, the most common modes of mutations in rabies virus, should not permit the transcription and translation of L as a protein separate from P. We passaged the RVdG-P-HCVP-L virus in producer cell lines for 8 months in the absence of ASV, periodically assaying viral sensitivity to ASV. This long term passaging did not detectably diminish ASV sensitivity (Fig. 5c), suggesting that this “cut-out” design can curb the emergence of escape mutants and maintain pharmaceutical regulation of the viral vector over a substantially more extended period of time than the previous design.

**Figure 5.**
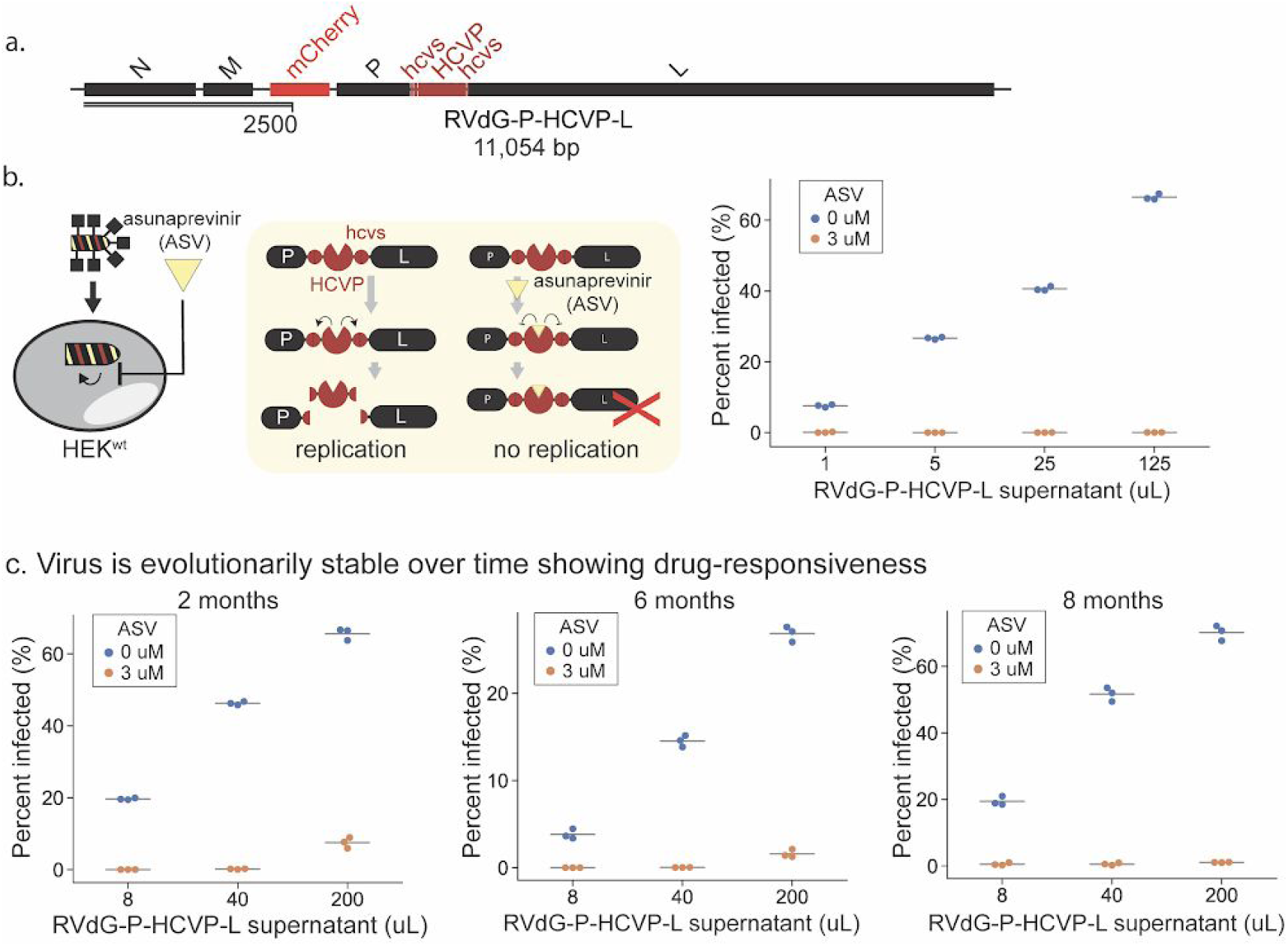
An external drug inhibits viral replication and permits viral removal. **(a)** Recombinant rabies genome sensitive to drug inhibition, RVdG-P-HCVP-L. **(b)** Asunaprevir (ASV) inhibits viral replication. Left, a Hepatitis C Virus protease (HCVP, red pac-man) flanked by its cleavage sites (hcvs, red circles) is inserted between the P and L proteins to create a P-HCVP-L fusion. HCVP cleavage of the flanking cleavage sites will separate P and L to permit viral replication. Addition of ASV, an HCVP inhibitor, will result in non-functional P-HCVP-L fusion proteins and inhibit RVdG-P-HCVP-L viral replication. **(c)** RVdG-P-HCVP-L exhibits evolutionary stability. RVdG-P-HCVP-L collected from 2, 6, and 8 months of continuous passage were tested on HEK^wt^. Regulation by ASV was maintained.

So far, we have focused on controlling the establishment of infection. However, for many applications, it will be crucial to terminate a productive infection after the viral vector and the encoded circuit perform their function. Therefore, we tested the ability of ASV to “cure” an established infection by the engineered virus. We used time-lapse microscopy to follow the dynamics of infection in the same population of cells over time before and after ASV addition (Fig. 6a).

**Figure 6.**
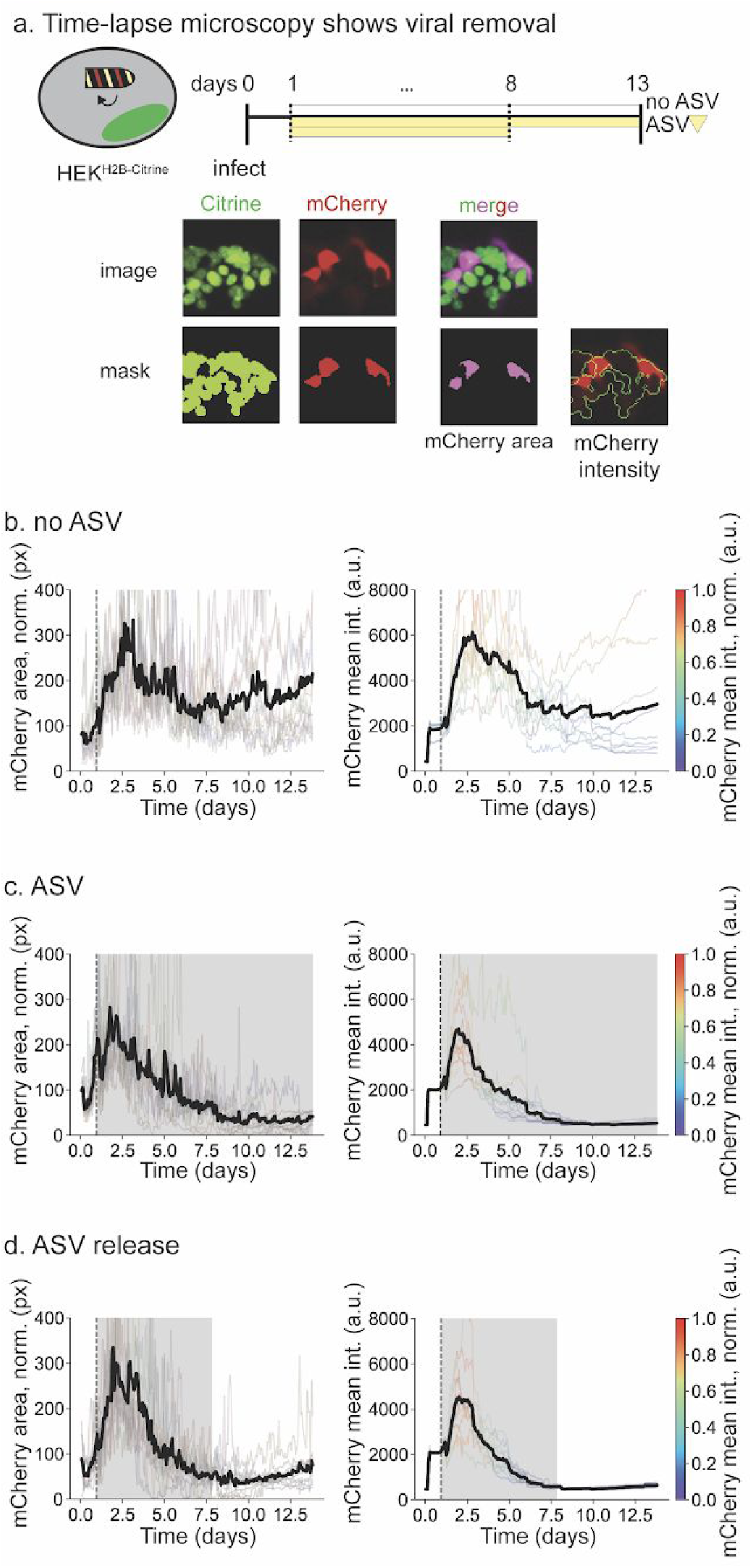
Time-lapse microscopy shows viral removal from established infections. **(a)** Top: Timeline of infection with no ASV (top blank bar), continuous ASV (middle yellow bar), or ASV-then-release (bottom shorter yellow bar). Bottom: Example of image processing. A binary mask was created based on the signal in the Citrine channel. The mCherry area shows the overlay between the mCherry mask and Citrine mask. Note that not all mCherry+ areas are Citrine+, because the tagged H2B-Citrine is localized in the nucleus while mCherry doesn’t have a localization tag. The mean intensity of mCherry signal in the Citrine mask was also quantified. HEK target cells expressing H2B-Citrine were infected for 24 hours with RVdG-P-HCVP-L (dotted line) and then cultured in media containing no ASV (**b**, no ASV), continuous ASV (**c**, ASV), or ASV for 6.9 days and then for 5 days in media containing no ASV (**d**, ASV release). Grey shading indicates the presence of ASV in media. Transparent and black lines respectively represent traces from individual movies and the mean of those traces. Fraction of infected cells as indicated by “mCherry area, norm (px)” (b, c, d, left column) was calculated as the fraction of mCherry+ pixels within the H2B-Citrine+ mask. Individual traces for mCherry intensity are color-scaled from purple to red, where mCherry intensity values are normalized by the maximum mCherry intensity within each trace. The slight increases at late times in the mean mCherry intensity traces represent cellular autofluorescence. In order to display all mean traces on the same y-scale, the tops of some individual traces are cut off. For complete traces, see Supplementary Fig. 3.

We first incubated HEK293 cells that express a constitutive H2B-Citrine (HEK^H2B-Citrine^) with viral particles, and quantified mCherry signal as a surrogate for the level of infection using two similarly behaving metrics (Fig. 6a, see Methods for details). After one day, we observed increased mCherry signal (Fig. 6b-d), consistent with viral infection and replication. This signal remained high for more than ∼13 days (Fig. 6b, Supplementary Movie 1, left). In a parallel experiment, we added ASV to the media after one day (Fig. 6c). This treatment led to a decay in the mCherry signal over the following 7 days, until no signal could be detected above background (Fig. 6c, Supplementary Movie 1, center). Thus, ASV addition successfully terminated viral protein expression.

Nevertheless, in principle low levels of virus could remain after ASV treatment and resume replication after ASV removal. We therefore asked whether transient addition of ASV could permanently cure an established infection of the engineered virus. First, in a preliminary experiment, we treated infected cells with two concentrations of ASV for 6.2 days (see Methods) and then removed the drug for 3 days. We observed no viral re-emergence in all but one field of view. This result suggested the potential for further improvement through repeated and prolonged ASV dosing (Supplementary Fig. 4a and 4b, Supplementary Movie 2). We therefore extended the period of ASV exposure to 6.9 days, during which we carried out daily media changes. We then washed out the drug, and continued to monitor the culture for 5 more days (Fig. 6d, Supplementary Movie 1, right), which is more than sufficient time for intracellular amplification. We observed no re-emergence of infection during this time period across all ten fields of cells (Fig. 6d). These results suggest that infections of the engineered virus could be successfully reversed by transient ASV addition in the overwhelming majority of cells.

## Discussion

Because they remain outside the nucleus and thereby avoid the potential risk for insertional mutagenesis, RNA viruses are attractive candidates for future synthetic circuit delivery vectors. In fact, while DNA vectors currently remain more prevalent in therapeutic applications, RNA viruses have received growing attention^40^, including from synthetic biologists seeking to improve their specificity ^41–43^. Here we sought to address key challenges required to make RNA viruses into a more engineerable, and safer, alternative to DNA vectors. Rabies virus, with its extensive history of engineering^44^ and applications in neuroscience ^19,20^, provides an ideal model system. As a step towards increasing the engineerabiity of rabies virus, this work had the dual purposes of integrating well-known mechanisms such as pseudotyping into a single platform, while also designing new mechanisms to address outstanding challenges such as evolutionarily stable drug control of replication. The modules analyzed here are for the most part not rabies-specific and likely to be transferable to other RNA viruses. We can now achieve multiple levels of specificity and control in rabies virus (Fig. 1). These levels include controlling viral secretion from “sender” cells (Fig. 2, 3c); achieving selective infection of target cells based on surface proteins, with external control through bispecific bridge proteins (Fig. 3); implementing conditional replication in target cells based on an intracellular protein (Fig. 4); and designing evolutionarily robust control of viral replication with the drug ASV (Fig. 5, Fig. 6). Because RNA viruses have elevated mutation rates compared to many DNA viruses ^34,45^, this last feature addresses a critical hurdle that will be necessary for RNA viral vectors to compete with DNA vectors in biomedical applications.

As a tool for basic research in neurobiology, the engineered rabies vectors introduced here could be immediately applicable for tracing synaptic connections with better control and higher specificity. Upon further optimization, these vectors might help realize a central biomedical promise of synthetic biology: they could be delivered in sender cells that naturally home to a disease tissue ^46^, exit sender cells only in the correct microenvironment, selectively enter target cells that display certain surface markers, replicate specifically in cells positive for specific markers and pathway activities, deliver genetic cargo to change cellular behavior as a therapy, and finally be eliminated with a small-molecule drug. The self-replication of such vectors guarantees that their cargo will be highly expressed so as to effectively perform their functions, and the RNA nature of their genome and the ability to eliminate the virus from infected cells minimize the risk of a synthetic circuit leaving “scars” in the host genome or intracellular signaling pathways. While many challenges to this vision undoubtedly remain, the results here provide proof of principle demonstration of core capabilities.

Beyond the artificial signals such as dox and GFP that control the current version of rabies vectors, one next step is to apply a similar design principle to engineering vectors whose replication is conditioned upon the activity of endogenous pathways and antigens, starting with those that drive oncogenesis^47,48^. For example, a virus that could conditionally replicate in cells with elevated Myc or Ras activity could provide enhanced selectivity for tumor cells.

In addition, to realize the potential of RNA viral vectors, we will still need to address several challenges. First, overly high vector replication and cargo expression in host cells after delivery could generate toxicity by competing for cellular resources, leading to detrimental effects on the patient. To address this, synthetic negative feedback circuits could be added to maintain vector/cargo levels below a tolerance threshold. Second, although rabies virus has evolved to counter detection and clearance by the immune system^49^, it, like most other RNA viruses, is nevertheless immunogenic. One solution would be to encode additional immunomodulating molecules in the vector, so that innate immunity can be temporarily suppressed while the vector and its cargo carry out their function. Third, although rabies vectors with inserts up to 3.7 kb long have been successfully produced and used for in vivo tracing^50^, it will be critical to systematically determine how delivery efficiency varies with the size and sequence identity of genetic cargo. If size limits prove to be inadequate for common applications, one could seek to identify related viruses with larger capacity or develop strategies to split cargo among two or more viruses, each of whose replication depends on the other.

Future optimization of RNA viral vectors would benefit from a plethora of novel methods to regulate RNAs and proteins, including the use of synthetic riboswitches^43^. These methods would provide additional means for controlling vector replication. On the other hand, the ability to bypass transcriptional control and host genome integration means that synthetic circuits made of RNAs and proteins are ideal cargo for RNA vectors, especially protein circuits that can be compactly encoded on a single transcript. We anticipate that engineering rabies viral vectors and their cargo could drive the development of other homologous engineered RNA viruses, each adapted to specific applications.

## Supporting information

Supplementary Figures 1, 2, 3, 4. Captions for Supplementary Movie 1, Supplementary Movie 2, Supplementary Table 1, Supplementary Table 2.

## Acknowledgements

We thank J. Ruan and A. Erskine for technical assistance; R. Zhu, Z. Singer, H. McBride, and L. Luo for critical feedback.

## Author contributions

X.J.G. and L.S.C. conceived of the project. X.J.G., L.S.C., M.S.K., and M.B.E. designed experiments. X.J.G., L.S.C., M.H.I. and M.S.K. performed experiments. X.J.G., L.S.C. and M.B.E. analyzed data. X.J.G., L.S.C., and M.B.E. wrote the manuscript, with input from all authors.

## Data and materials availability

All DNA constructs will be available from Addgene, and cell lines are available from M.B.E. under a material transfer agreement with Caltech. The datasets generated and analyzed and the computer code used during the current study are available at data.caltech.edu, DOI 10.22002/D1.1438. Flow cytometry data analysis software used for this study is available at https://antebilab.github.io/easyflow/.

## METHODS

### Plasmid construction

All constructs were generated using standard procedures. The backbones were linearized using restriction digestion or PCR, and inserts were generated using PCR or gBlock synthesis (IDT). A list of all plasmids reported in this manuscript is included in Supplementary Table 1, and all sequences were deposited to Addgene.

### Tissue culture

Flp-In™ T-REx™ 293 Cell Line (Human Embryonic Kidney cells that contain a single stably integrated FRT site at a transcriptionally active genomic locus, and stably expressing the tetracycline repressor protein) were purchased from Thermo Fisher Scientific (R78007). B7GG cell line (Baby Hamster Kidney cells that contain a stably integrated T7 RNA polymerase and rabies glycoprotein) and HEK-TVA cell line (Human Embryonic Kidney cells that contain stably integrated TVA receptor) were kindly gifted from Dr. Lindsay Schwarz at St. Jude Children’s Research Hospital.

HEK293 derived cells were cultured in a humidity controlled chamber at 37°C with 5% CO_2_ in media containing DMEM (Thermo Fisher Scientific, cat no. 11960-069) supplemented with 10% FBS (VWR, 76308-946), 1 mM sodium pyruvate (Thermo Fisher Scientific, cat no. 11360-070), 1 unit/ml penicillin, 1 μg/ml streptomycin, 2 mM L-glutamine (Thermo Fisher Scientific, cat no. 10378-016) and 1X MEM non-essential amino acids (Thermo Fisher Scientific, cat no. 11140-050) (293 media). B7GG derived cells producing rabies virus were cultured in a humidity controlled chamber at 37°C with 5% CO_2_ in media containing DMEM supplemented with 2.5-5% FBS, 1 mM sodium pyruvate, 1 unit/ml penicillin, 1 μg/ml streptomycin, 2 mM L-glutamine and 1X MEM non-essential amino acids (BHK media).

100 ng/mL doxycycline was added whenever expression is needed from a CMV/TO promoter. Trimethoprim (TMP) was delivered at 1 µM. Asunaprevinir (ASV) was delivered at 3 µM.

### Cell line construction

To generate cell lines with stably integrated transgenes antibiotic selection was performed. Flp-In™ T-REx™ 293 Cell Line or BHK21 cells were transfected in 24-well plates and transferred two days later into a 6-well plate containing selection media (Supplementary Table 2). After PiggyBac-based integration, monoclonal cell populations were established through limiting dilution, and preliminary screening was performed to identify clones with highest transgene expression using flow cytometry (Supplementary Table 2). Transgenesis using pOG44 Flp-recombinase into Flp-In™ T-REx™ 293 Cell Line resulted in singly-integrated cell lines.

### Production of Packaging Cell Lines

B7GG was integrated with a transgene that encodes for constitutive co-expression of TEVP and Cerulean (B7GG-TCer). This packaging line was generated to permit efficient production of modified rabies genomes containing an essential protein tagged with a destabilizing domain, in which a TEVP cleavage site was inserted between the essential protein and the degron. TEVP cleavage of the degron stabilizes the essential protein allowing viral replication.

### Production of Rabies Viruses

Viruses were initially established using the protocol as described in Osakada, Nat Pro. 2013 and scaled for 24-well plates. RVdG-P-DHFR and RVdG-P-GBP were produced in B7GG-TCer producer lines while RVdG and RVdG-P-HCVP-L viruses were produced in either cell line (Supplementary Table 2). Briefly, B7GG and B7GG-TCer cells were plated in 24-well plates at 0.05×10^6^ cells per well in 293 media. Cells were transfected with a DNA mixture containing 240 ng rabies genome, 120 ng pcDNA-SADB19N, 60 ng pcDNA-SADB19P, 60 ng pcDNA-SADB19L and 40 ng pcDNA-SADB19G. Media was changed one day after transfection to BHK media. Cells were transferred from a 24-well plate to a 6-well plate three days after transfection and maintained in BHK media. Fresh BHK media was added every day to facilitate viral spread. Viral spread was checked every day using fluorescent microscopy. Supernatant collection only proceeded when the virus had visibly infected the majority of the population.

Amplified rabies viruses were concentrated and purified as described in (Osakada, Nat Prot. 2013) to generate starter viral stocks. Briefly, infected producer lines were expanded to three 15 cm dishes and exchanged with fresh BHK media the following day. After three days, conditioned media was collected and filtered through at 0.45 um filter. The supernatant was concentrated in an ultracentrifuge at 70,000 g for 2 hr at 4 °C on a 20% sucrose pad. Viral pellets were resuspended in 1X Hank’s Balanced Salt Solution (HBSS) and stored at -80 °C for future use.

Pseudotyped EnvA rabies virus was purchased from Janelia Farms.

### Production of Bridge Protein Cell Lines

#### In vivo production of bridge proteins

The extracellular domain of ASLV-A envelope protein from the plasmid pAAV-TRE-HTG (Addgene) and the targeting domain encoding the Gbp6 nanobody^33^ were combined by PCR and cloned into a Piggybac transfer vector under a synthetic constitutive promoter (Supplementary Table 1). Flp-In™ T-REx™ 293 cells were stably integrated to create a polyclonal bridge protein secretion line. To collect produced bridge proteins, the stable line was seeded at a density of 0.1×10^6^ cells per well of a 6-well plate and cultured under standard conditions overnight. The media was exchanged the following day for BHK21 media and cells were cultured over two days with additional fresh media applied the second day. Conditioned media was collected and centrifuged at 13,000 rpm for 20 minutes to remove cellular debris. 300 uL of conditioned media was applied in each condition unless otherwise noted.

#### In vitro production of bridge proteins

Bridge proteins were purified at the Caltech Protein Expression Center. Briefly, 6XHIS-tagged bridge proteins encoding plasmids (Supplementary Table 1) were transfected into Expi293FTM cells and conditioned media was harvested four days post-transfection. The supernatant was spun at 2000 rpm for 10 minutes and proteins were purified with Ni-NTA affinity chromatography (Cytiva, 5 mL HisTrap). Expression and purification were checked by SDS-PAGE and Coomassie Blue staining. Protein concentration was quantified using PierceTM 660 nm (Thermo Fischer, cat no. 22662) assay.

### Transient Transfection

293 cells were seeded at a density of 0.05×10^6^ cells per well of a 24-well plate and cultured under standard conditions overnight. The following day, the cells were transfected with plasmid constructs using Lipofectamine LTX (Thermo Fisher) as per manufacturer’s protocol.

### Infection with Rabies

Infected producer cell lines B7GG and B7GGTCer were plated to 60% confluency in 6-well plates. The following day, the media was changed to 2 mLs of fresh BHK media. Two days later an additional 1 mL of BHK media was added. Virus containing media was collected on the third day and supernatant was spun down for 5 minutes at 15,000 rpm to remove cellular debris. Supernatant from these freshly prepared stocks were used directly in RVdG-P-GBP (Fig. 4) and RVdG-P-HCVP-L (Fig. 5, Fig 6, Supplementary Fig. 3, Supplementary Fig. 4) experiments. Viral amount was determined by volume, as indicated on figures. Infection using purified RVdG-EnvA was performed by calculating MOI based on the reported viral titer provided by Janelia Farms.

For experiments containing sender cells (Fig. 2, Fig. 3c), sender cells were plated at 30% confluency and the following day media was exchanged with 1 mL BHK media and 1 mL of freshly prepared RVdG supernatant, as described above.

All experiments were conducted in 24-well plates. In co-culture experiments, 30,000-45,000 cells for each target population were counted and pre-mixed prior to plating. Specifically, for the co-culture performed in Fig. 4b, the expression of Citrine was induced with doxycycline overnight prior to plating and was present throughout the experiment (Supplementary Table 2, refer to HEK^doxCit^). For single target cell population experiments, the cells were seeded at 75,000 cells per well. Infected sender cells, virus-containing supernatant, purified virus, bridge protein, and compounds were added to each well immediately after target cells were plated unless otherwise noted.

### Flow cytometry

Three days after infection, cells were prepared for flow cytometry by trypsinizing with 40 uL of 0.05% trypsin for 1 min at room temperature. Protease activity was neutralized by resuspending the cells in buffer containing 100 uL of HBSS with 2.5mg/ml Bovine Serum Albumin (BSA) and 1 mM EDTA (Thermo Fischer Scientific, cat no. 15575020). Cells were then filtered through 40 μm Falcon™ Cell Strainers (Thermo Fischer Scientific, cat no. 08-771-1) and analyzed by flow cytometry (MACSQuant VYB, Miltenyi or CytoFLEX, Beckman Coulter). Matlab-based software for processing flow cytometry data was developed in-house by Yaron Antebi.

### Fluorescent signal quantification from flow cytometric measurements

Figures 2, 3, 4, 5 and Supplementary Figure 1 present quantitative analysis of flow cytometry measurements. For each sample in a comparison group (experiments performed in the same batch and data shown on the same plot), we reported the percent infected as identified by mCherry+ cells.

#### Co-culture gating

The fluorescence values of each cell line were determined in the constitutive fluorescent signal channel (Citrine or IFP in most cases). Each cell line was analyzed independently. For example, for a co-culture containing HEK^wt^ and HEK^Cit^, monoculture samples of HEK^wt^ and HEK^Cit^ were each analyzed independently in the Citrine fluorescent channel. For analysis, we fit the fluorescence log distribution with skew Gaussian distributions, i.e. *n* × normcdf(*x,m,k*) × normpdf(*x,m,s*) in Matlab using non-linear least-square fitting, and reported the mode of the resulting fit. Here, the normcdf(*x,m,k*) and normpdf(*x,m,s*) functions are cumulative probability density and probability density functions for Gaussian distributions, respectively. The parameter *n* is a normalization factor, the parameter *m* is the mean of each Gaussian distribution, *s* is the standard deviation of the probability density function that parameterizes the width of the distribution, and *k* is the standard deviation of the cumulative probability density function that parameterizes the skewness of the distribution.

#### Percent infected calculation

The mode of the resulting fit is used to gate different cell types from the co-cultured samples. For monoculture experiments, gating was not performed. For all cells within the gate in each sample, we fit a cubic spline and identified the local minima between the two peaks of mCherry+ and mCherry-cells. An average local minimum was calculated across all bimodal samples. We use this average local minima as a threshold between non-infected/weakly infected cells (mCherry-) and highly infected cells (mCherry+). A percent infected metric is calculated from the resulting histogram by calculating the area both above and below the threshold:

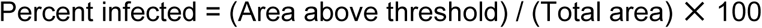

### Sequencing virus genomes P-DHFR and P-GBP

Three independent viral cultures were passaged for 2-4 months prior to genome sequencing. Reverse transcription of viral RNA was performed on samples using the Thermo Fisher Scientific protocol for SuperScript IV. PCR amplification of the genome using forward primer (5’-ACCCTCCAGGAAAGTCTTC-3’) and reverse primer 5’-AATAGGGTCATCATAGACCTCTC-3’) were gel purified and submitted for Sanger sequencing with Laragen Sequencing.

### Time-lapse microscopy

For time-lapse imaging of rabies dilution (Fig. 6, Supplementary Fig. 3, Supplementary Fig. 4, Supplementary Movies 1 and 2) 5,000 H2B-Citrine were plated on 24-well glass-bottom plates (CellVis). Cells were induced with 100 ng/mL doxycyline overnight in normal culturing conditions. The following morning, the media was replaced with imaging media containing FluoroBrite DMEM (Thermo Fisher) supplemented with 10% FBS, 1 mM sodium pyruvate, 1 unit/ml penicillin, 1 μg/ml streptomycin, 2 mM L-glutamine and 1X MEM non-essential amino acids and 100 ng/mL doxycycline. All time-lapse images were acquired on an inverted Olympus IX81 fluorescence microscope with Zero Drift Control (ZDC), an ASI 2000XY automated stage, iKon-M CCD camera (Andor, Belfast, NIR), and a 20X PAN-FL objective (1.42 NA). Fluorophores were excited with an X-Cite XLED1 light source (Lumen Dynamics). Cells were kept in an environmental stage-top chamber enclosing the microscope, with humidified 5% CO_2_ flowing at 37°C (Okolab H301-K stage top with an O_2_-CO_2_ UNIT-BL mixer). Microscope and image acquisition were controlled by Metamorph software version 7.10 (Molecular Devices).

Imaging started approximately 2 hours after changing the media to fluorescent imaging media. B7GG lines producing virus were cultured for three days in imaging media. Conditioned media was collected and centrifuged at 13,000 rpm to pellet cellular debris. 250 uL of the centrifuged supernatant containing virus was added after approximately 2 hours of imaging to infect cells. The initial infection was established for 24 hours. To observe the removal of the rabies genome with prolonged ASV treatment, cells were washed with imaging media three times to remove residual virus and then incubated with imaging media containing 3 μM of ASV. Media was changed every 24 hours. To confirm that the virus does not re-emerge after ASV removal, the ASV-containing imaging media was replaced with normal imaging media after 7 days.

Images were acquired every 90 minutes throughout the duration of the movie. Cells that were in the field of view before infection and remained alive and visible in the field of view without death for at least five days were used for initial data analysis through manual inspection.

In the preliminary experiment (Supplementary Fig. 4, Supplementary Movie 2), cells were infected for 17 hours and media was changed every 3 days over 6 days. Images were acquired every 60 minutes. For analysis, a similar selection criteria was performed in which cells that were infected and remained within the field of view for five days were used for data analysis. Additionally, we excluded from analysis fields of view that were out of focus, near the edge of the well, or exhibited clumping or apoptosis within the last day of imaging.

### Movie analysis

#### Data processing

mCherry mean intensity values were calculated based on total levels of fluorescence in the mCherry fluorescent channel as identified by the position of cells in the Citrine fluorescent channel. To systematically identify the position of cells, total constitutive Citrine signal was used to segment each image. The resulting segmentation mask was used to calculate the number of mCherry+ pixels within each region. To capture the magnitude of rabies expression within infected cells, the mean intensity was calculated for each mCherry+ region and averaged for each timepoint. The fraction infected metric was calculated by identifying the mCherry+ areas within the Citrine+ areas.

We first estimated the fraction of cells infected by rabies virus. We used the constitutive Citrine signal to generate a mask for all cells, where a pixel is considered to belong to a cell if its Citrine intensity is > 400, as determined by an Otsu threshold of the first Citrine containing image. Within the Citrine mask, pixels are considered rabies-positive when mCherry intensity is greater than a moving Otsu threshold to account for fluctuations in background signal. As such rabies-positive area divided by the area of the Citrine mask (mCherry area norm. in Fig. 6, Supplementary Fig. 3 and Supplementary Fig. 4) is reported as a proxy for the fraction of rabies-infected cells in each image. The constitutive Citrine signal is localized to the nucleus and may underestimate the rabies-positive area, especially towards the beginning of the imaging when rabies-encoded mCherry form puncta in the cytoplasm.

In addition to thresholding mCherry signal, we also estimated the quantitative level of rabies infection by calculating the total mCherry intensity within each Citrine positive region of each image and averaged across these regions (mCherry mean int. in Fig. 6, Supplementary Fig. 3 and Supplementary Fig. 4) as a proxy for the level of rabies infection in that image.

#### Movie generation

For visualization purposes, the timepoint at which maximum mCherry intensity occurs for each movie was determined and used to rescale all images between the 5th and 99.5th percentile intensity values.

### Data and code availability

The datasets generated and analyzed and the computer code used during the current study are available at data.caltech.edu, DOI 10.22002/D1.1438. Flow cytometry data analysis software used for this study is available at https://antebilab.github.io/easyflow/.

