## Supplementary Figures 1, 2, 3, 4. Captions for Supplementary Movie 1, Supplementary Movie 2, Supplementary Table 1, Supplementary Table 2. for "Engineering multiple levels of specificity in an RNA viral vector"

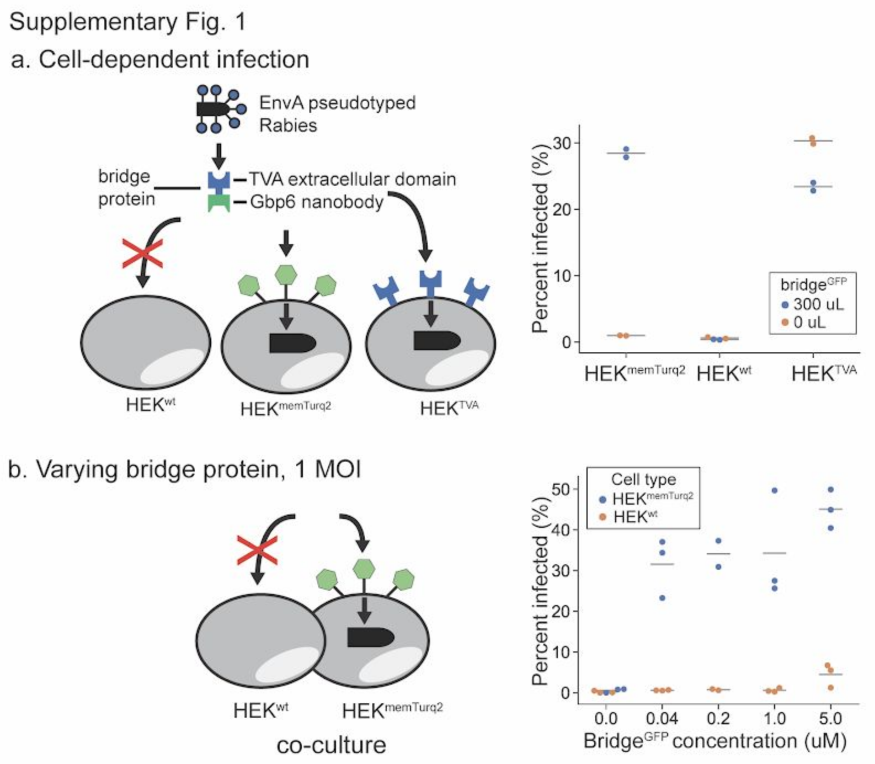


**Supplementary Figure 1: Pseudotyped rabies viral entry is dependent on presence of bridge and cell-surface marker. (a)** RVdG-EnvA virus infection of independently cultured $HEK^{memTurq2}$, $HEK^{wt}$, and $HEK^{TVA}$ with or without $bridge^{GFP}$. **(b)** RVdG-EnvA and varied $bridge^{GFP}$ concentrations infection of a co-culture of $HEK^{memTurq2}$ and $HEK^{wt}$.

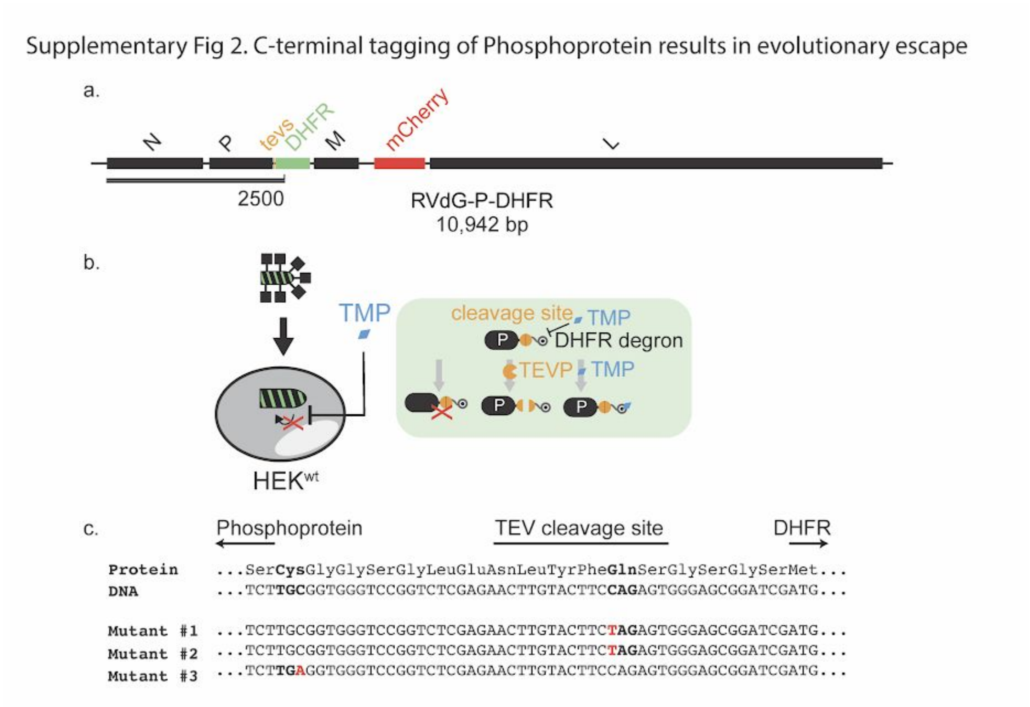


**Supplementary Figure 2: C-terminal tagging of Phosphoprotein exhibits evolutionary escape.** **(a)** Recombinant rabies genome, RVdG-P-DHFR. **(b)** Schematic detailing phosphoprotein regulation through C-terminal DHFR degron tagging. The phosphoprotein is stabilized by removal of the DHFR degron either through TEVP cleavage (orange pac-man) of the corresponding cleavage site (orange circle)**.** Trimethroprim (TMP, blue diamond) inhibits the degron and stabilizes the reporter. **(c)** Sequencing of three escape mutants identified two distinct missense mutations. Coding sequences and corresponding amino acid sequences of Phosphoprotein, TEVP cleavage site, and DHFR are indicated above. Red indicates single nucleotide mutations and bold indicates mutated codons.

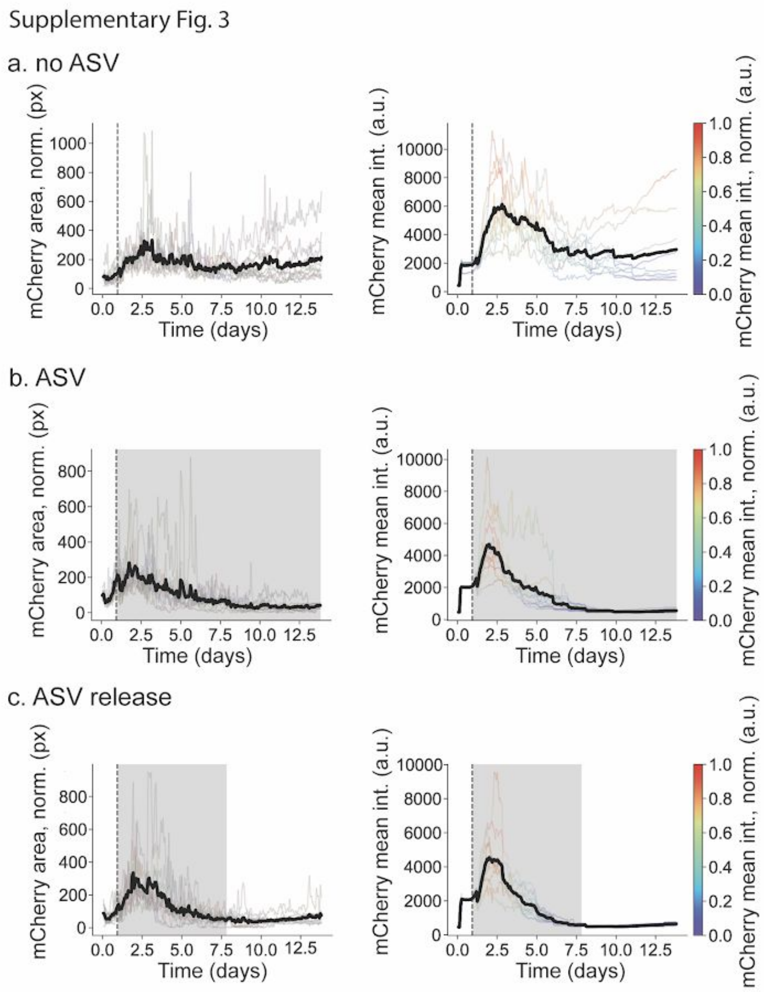


**Supplementary Figure 3: Time-lapse microscopy shows viral removal of non-infected cells.** Plots shown from Figure 5d with full y-axis. HEK target cells expressing H2B-Citrine were infected for 24 hours with RVdG-P-HCVP-L (dotted line) and then cultured in media containing no ASV (**b,** no ASV), continuous ASV (**c,** ASV), or ASV for 6.9 days and then for 5 days in media containing no ASV (**d,** ASV release). Grey shading indicates the presence of ASV in media. Transparent and black lines respectively represent traces from individual movies and the mean of those traces. Fraction of infected cells as indicated by “mCherry area, norm (px)” (b, c, d, left column) was calculated as the fraction of mCherry+ pixels within the H2B-Citrine+ mask. Individual traces for mCherry intensity are color-scaled from purple to red, where mCherry intensity values are normalized by the maximum mCherry intensity within each trace. The slight increases at late times in the mean mCherry intensity traces represent cellular autofluorescence.

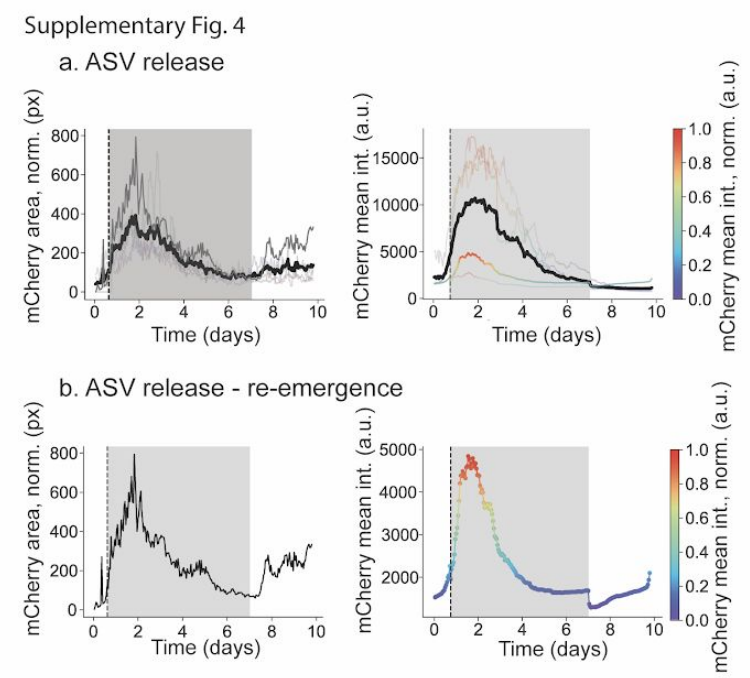


**Supplementary Figure 4: Time-lapse microscopy shows viral reemergence in a preliminary experiment. (a)** HEK target cells expressing H2B-Citrine were infected for one day with RVdG-P-HCVP-L (dotted line) and then cultured in media containing ASV for 6.2 days with media changed every three days and then for 3 days in media containing no ASV. Grey shading indicates the presence of ASV in media. Transparent and black lines respectively represent traces from individual movies and the mean of those traces. Opaque lines indicate viral re-emergence, shown independently in **(b)**.

**Supplementary Movie 1:** Example RVdG-P-HCVP-L infected cells with no ASV treatment (left), continuous ASV treatment (center) and ASV treatment for 6.9 days and then in media containing no ASV for 5 days (right). Top is the mCherry channel and bottom is the H2B-Citrine channel. Timestamp in the upper left corner of the mCherry channel in the first column indicates days:hours:minutes for all conditions. Yellow triangle in the upper right corner indicates ASV addition.

**Supplementary Movie 2:** Preliminary experiment in which example RVdG-P-HCVP-L infected cells with a single ASV treatment for 6.2 days and then in media containing no ASV for 3 days. Top is the mCherry channel and bottom is the H2B-Citrine channel. Timestamp in the upper left corner of the mCherry channel in the first column indicates days:hours:minutes for all conditions. Yellow triangle in the upper right corner indicates ASV addition. Re-emergence of the virus can be seen in the right movie.

**Supplementary Table 1:** List of constructs used to generate cell lines and rabies viruses.

**Supplementary Table 2:** List of cell lines used for experiments and cell lines used to produce rabies viruses and bridge proteins. Highlighted blue rows indicate purchased cell lines. Producer lines are used to package the rabies virus purified for infection. Bridge producing lines are used to generate bridge protein. Sender lines are integrated with a viral glycoprotein to transmit the virus to Receiver lines in the co-culture studies. See Methods for details.
